# Deciphering the genetic basis of phytoplankton traits through genome-wide association studies

**DOI:** 10.64898/2026.02.27.708454

**Authors:** Agathe Maupetit, Vincent Segura, Anne Pajot, Elodie Nicolau, Gael Bougaran, Thomas Lacour, Jean-Baptiste Berard, Aurelie Charrier, Nathalie Schreiber, Elise Robert, Bruno Saint-Jean, Grégory Carrier

## Abstract

While marine genomic databases are rapidly expanding through large-scale sequencing programs, the functional characterization of these genomes progresses slowly and remains a major scientific challenge. In this study, we applied a Genome-Wide Association Study (GWAS) approach to identify and investigate genomic loci without *a priori* assumptions.

The microalga *Tisochrysis lutea* was selected as a case study due to its economic importance and the extensive knowledge accumulated over the years. Particular attention was paid to pigment and lipid metabolism due to their high commercial value. To implement the GWAS approach, a collection of 100 algal lineages was established from 15 available polyclonal strains. This collection was then phenotyped under two different culture conditions. Of the 31 phenotypic traits investigated, 18 met the requirements for GWAS analysis. Concurrently, each algal lineage was genotyped by whole-genome sequencing to inventory all genetic polymorphisms. A mixed model was applied, revealing 13 significant associations between phenotypic traits and alleles. These associations highlight previously unsuspected genomic loci that play a major role in pigment or lipid content.

The candidate genes identified at these loci may have a direct or indirect role in these metabolic pathways. Nevertheless, elucidating the precise molecular mechanisms of the associated genes remains limited without the implementation of functional approaches. Despite the complexity of the process, we conclude that the GWAS approach is an effective strategy for exploring phytoplankton genomes, particularly for quantitative traits of interest. Ideally, this approach should be combined with other functional methods to progressively characterize marine genomes.

## Introduction

Following the sequencing revolution, the subsequent scientific challenge is no longer just generating “big data,” but rather deciphering the functional roles of these genes (Chen, 2025). For instance, a reference collection of 116 million genes from global plankton communities was established, data showed that while approximately 40% possessed orthologues with genes or proteins of known function, three-quarters still lacked a defined molecular function (Carradec et al., 2018; Salazar et al., 2019). This bottleneck in functional annotation remains significant for all species, but it is even greater for marine organisms, which are notoriously poorly characterized. Consequently, relying solely on sequence homology to determine protein or gene function is often insufficient in the marine species (Cleves et al., 2020; Rumin et al., 2020).

The traditional approach to characterizing gene function involved the generation of mutants to analyse the resulting phenotypes. For example, mutant libraries have been created in the model microalgal *Chlamydomonas reinhardtii* to identify genes involved in photosynthesis or heat tolerance (Li et al., 2019; Mattoon et al., 2023). However, creating these libraries requires considerable resources. Although CRISPR-based tools have improved the efficiency of mutagenesis, stable genetic transformation has only been mastered in a limited number of marine organisms (Mochdia and Tamaki, 2021; Przybyla and Gilbert, 2022). These methods are efficient but they work “gene by gene” and are thus very time-consuming, moreover they require mastered genetic transformation that is not easy in marine organism. More recently, integrative multi-omics and machine learning approaches have emerged in human medicine and in a few model organisms. However, their application to marine organisms remains limited due to a lack of data and knowledge for identifying robust candidate genes (Dobretsov and Rittschof, 2023; Kwon et al., 2023; Williams, 2024).In this work, we propose to use Genome-Wide Association Study (GWAS) to identify several candidate genes involved in key biological processes (Uffelmann et al., 2021). The aim of GWAS is to identify statistical associations between DNA polymorphisms and quantitative traits (Korte and Farlow, 2013). GWAS has successfully been applied to identify genes involved in human diseases (Prata et al., 2019; Visscher et al., 2017), livestock breeding traits (Raza et al., 2020; Sharma et al., 2015), agricultural or ecological traits in plants (Alqudah et al., 2020; Bazakos et al., 2017; Hamdani et al., 2019). However, its application in marine organism remains limited so far.

Among the diversity of oceanic phytoplankton species and our limited knowledge about them, choosing a relevant species for a GWAS approach is not easy. In this study, we applied GWAS to advance our understanding of marine genomics using the golden-brown microalga *Tisochrysis lutea* as a model. This species was selected for two main reasons: firstly, its strong potential in aquaculture due to its content in secondary metabolites of interest (García-García et al., 2024); and secondly, because its cultivation has been well-mastered in our laboratory for several years (Bougaran et al., 2012). This species represents an intermediate level of knowledge, reflecting our current understanding of the marine environment. While it has been well described and basic genomic and physiological knowledge is available, this species lacks the extensive resources developed for reference microalgal models like *Chlamydomonas reinhardtii*. Historically, *T. lutea* was widely used as a food source in aquaculture since the 1970s because it was appetising and could be easily cultivated in hatcheries (Guedes and Malcata, 2012; Kaplan et al., 1986). Various studies were subsequently conducted on its biology, ecophysiology, genomics, and metabolism (Berthelier et al., 2023; Carrier et al., 2024; Nef et al., 2020; Pajot et al., 2023). However, fundamental biological knowledge regarding *T. lutea* remains limited, particularly concerning its ploidy and life cycle, which are currently uncharacterized. We hypothesize that it possesses a haplo-diplontic life cycle, similar to the well-studied species *Emiliania huxleyi*. However, despite several attempts, we have not yet succeeded in synchronizing its cycle in culture. Consequently, for the purpose of this study, we consider these microalgal cultures as an asynchronous population comprising individuals that may be either haploid or diploid.

Selecting phenotypic traits of interest was the second step in designing the GWAS approach. In most cases, the traits considered are those with human interest (economic, health, etc.). Thanks to its high antioxidant content, *T. lutea* is today of great interest for biotechnology applications (Bigagli et al., 2021; Chatzopoulou et al., 2024; Sathasivam and Ki, 2018). Lipids and pigment molecules are high-value compounds beginning to be exploited in this species (Bigagli et al., 2021; Chatzopoulou et al., 2024; Sathasivam and Ki, 2018). Specifically, it produces secondary pigments such as fucoxanthin, which was exploited for its protective role against cancer, inflammatory diseases and obesity (Liu et al., 2020; Miyashita et al., 2020). *T. lutea* also contains lipids with high added-value, including docosahexaenoic acid (DHA). This was an essential nutrient for brain development and function, and more than 90% of the world’s population has an insufficient supply of it. (Bazinet et al., 2020; Jovanovic et al., 2021; Madore et al., 2020). Since the 1970s, the cultivation conditions were established and progressively optimized (Kaplan et al., 1986). Recently, domestication programs were carried out to optimise lipid and pigment contents (Bougaran et al., 2012; Gachelin et al., 2021). However, the biological roles and the genetic basis of the biosynthesis of these compounds are only partially described, which is precisely why this study focused on these traits.

Based on decades of ecophysiological and genomic research and its growing interest in biotechnology sector, *T. lutea* represented an ideal model for conducting GWAS to interest genetic traits. In this study, the objective was to demonstrate the efficiency of GWAS as a tool for genomic characterization. We focused on pigment and lipid biosynthesis due to their significant industrial and economical value. The first challenge was to build a robust dataset, including genome-wide polymorphisms and quantitative traits on a set of algal lineages. In this aim, (1) we built up a collection of 100 *T. lutea* algal lineages; (2) we conducted more than 400 algae culture experiments to precisely phenotype individual algal lineages, measuring their pigment and lipid profiles; (3) we simultaneously sequenced the entire genome of each algal lineage. Finally, (4) a GWAS was performed to identify polymorphisms significantly associated with phenotypic variation, thereby unravelling the genetic architecture underlying pigment and lipid production in this microalga.

## Results and analysis

A wide range of results were measured during this experiment, and all data have been made available on GitLab (https://gitlab.ifremer.fr/gc0ca62/dynalgue-project), particularly the supplementary tables and figures.

### Collection of individuals set up for GWAS

In order to undertake GWAS, a large set of individuals lacking close ties of kinship was required (*i.e* at least 50 individuals) (Ott et al., 2011). However, the availability of biological resource is often limited for marine organisms. Concerning *T. lutea*, only 15 parental strains from different marine locations were available (Table S1.16). To address this issue, albeit imperfectly, we proposed exploiting the individual diversity within these parental strains available. Indeed, microalgal strains are generally not composed of pure lines, but rather encompass a broad range of genotypes (Lakeman et al., 2009; Sjöqvist, 2022). This intraspecific diversity within these parental strains was described in a previous study (Carrier et al., 2024). To establish a collection of 100 algal lineages, single cells were isolated from 15 parental strains by sorting their lipid and pigment profiles with cytometry (Sup. data S1.0 at S1.15). Each single cell was cultivated to give rise to an algal lineage. This approach was done to increase the probability of obtaining algal lineage with different lipid and pigment profiles. We hypothesized that the algal lineages from this collection would be sufficiently diverse, both genetically and phenotypically, to constitute an adequate collection for a GWAS.

### Phenotypic data for GWAS analysis

Quantitative analysis is based on the fundamental principle that an individual’s observed phenotype (*P_i_*) is a function of: its genotype (*G_i_*), the environment in which the phenotype is measured (*E*), and the interaction between the environment and the genotype (*P_i_ = G_i_ + E + G_i_. E*) (Falconer, 1996; Fisher, 1919). Therefore, in order to measure the correlation between phenotypic and genotypic variations, it was essential to control environmental variations as much as possible. In our study, to minimize the effect of environmental variations, the 100 algal lineages were phenotyped under highly controlled conditions, and sampled under the same physiological conditions using a dedicated culture device (Sup. data S2.1.1). Furthermore, to observe phenotype-genotype correlations that are not contingent upon environmental variations, it is required to perform phenotype measurements under various environmental conditions. Thus, experiments were conducted with two nutrient limited media: nitrogen-limited (Nlim) and phosphorus-limited (Plim) conditions. Phenotyping of each algal lineage was conducted when nutrient limitation was observed (Sup. figures S2.1.2 and S2.1.3). These nutrient limitations were applied to *T. lutea* cultures because they were commonly encountered by the microalgae in open ocean and they were known to have an impact on its phenotype and growth (da Costa et al., 2017; Huang et al., 2019). In these experimental conditions, a wide set of traits were measured for each algal lineage: physiological traits such as grow rate, (Sup. data S2.3), photosystem activity (Sup. data S2.2.4), carbon and nitrogen cells composition (Sup. data S2.2.2-S2.2.3); morphological traits such as cell size (Sup. data S2.1) and biochemical traits such as pigment profiles (Sup. data S2.4.3) and lipid profiles (Sup. data S2.5.1-S2.5.2).

#### Phenotypic variations of traits

One prerequisite to apply GWAS is that measured phenotypic variations between algal lineages were significant and, ideally highly genetically determined, so heritable. Several statistical analyses were performed on the 31 phenotype traits measured within each growing condition (Nlim and Plim) and for the 100 lineages to identify traits that can be analyzed in GWAS (Sup. data S3.1). First, variation of each trait within lineages was controlled by examining the interquartile range coefficient. For 19 traits in Nlim and 21 traits in Plim, range variation was sufficient to perform GWAS analysis (Sup. figure S3.2). Moreover, for each trait, their distribution was assessed, and according to a Kolmogorov-Smirnov test, 16 traits in Nlim and 18 traits in Plim condition approached a normal distribution (Sup. figure S3.3). Heritability (H²) was then estimated, and 24 traits had a value greater than 0.2, suggesting that their phenotypic variation was under genetic control (Sup. figure S3.5). Consequently, among the 31 phenotypic traits measured in each growing condition, only 18 met the prerequisites to investigate genetic-phenotypic associations (Sup. data S3.6).

#### Decomposition of phenotypic variance

As previously stated, effects of environment variables and history origins of individuals were linked to the phenotypic traits measured (Devlin and Roeder, 1999; Orgogozo et al., 2015). Therefore, variables as culture conditions effect (Nlim and Plim) and parental strains origin for each lineage (see collection set up, the previous paragraph) were retained. Thus, in order to estimate the phenotypic variance components for each measured trait, a mixed model was constructed, estimating each component of the variance (see materials and methods). Statistical tests indicated that the phenotypic variance of each trait was mainly attributable to the difference between parental strains and culture conditions. The importance of the parental strains component in the variance was confirmed by *Q*_ST_ values indices (Spitze, 1993), which were higher than 0.5 for most traits (Sup. table S3.8). In contrast, the part of variance attributable to differences between the algal lineages within each strain was limited. Moreover, the analyses revealed less variation between the different lineages within each parental strain than between the parental strains themselves (Sup. table S3.8). To observe and confirm the structure of variance and identify patterns of correlation among traits a principal component analysis (PCA) was performed on all measured traits across the 2 growing conditions (Figure 1). Culture conditions and parental strains were considered as supplementary variables to not influence axis construction. Their representations in PCA confirmed the results of previous model: the culture conditions exert a greater influence on traits surpassing the impact of parental strains. The physiological responses of algal cells appeared to differ significantly depending on whether phosphorus or nitrogen was limiting. These observations demonstrate that the effects of the two limitations on traits under the given culture conditions were unfortunately too significant and specific to be considered as a whole. Indeed, it has been well documented that nitrogen and phosphorus limitations induce specific physiological response in algae (Davies and Sleep, 1989; Rasdi and Qin, 2015), although we had not anticipated it would be to this degree.

**Figure 1.**
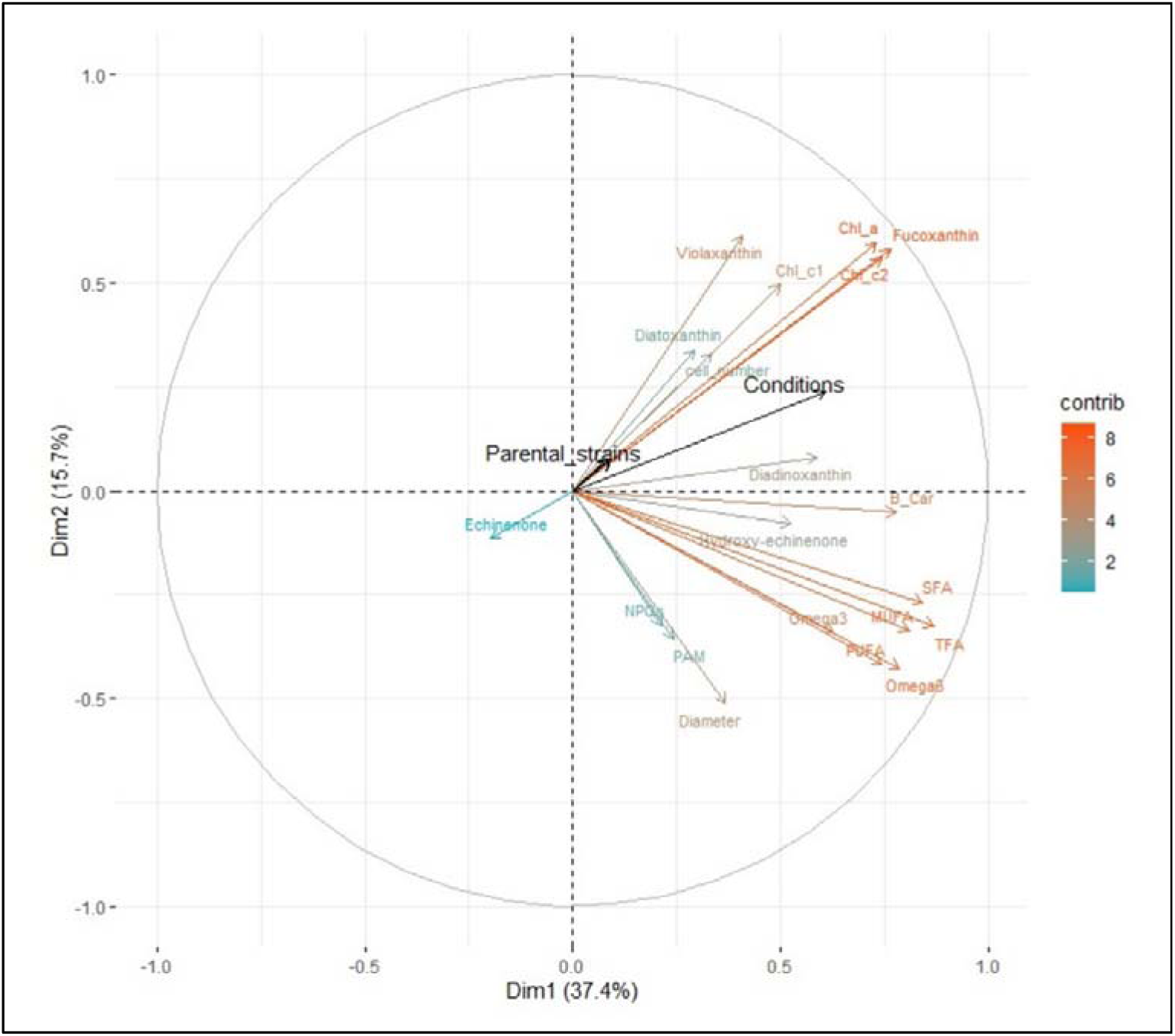
PCA based on phenotypic traits in both cultural conditions. *Each trait has been vectorized according to its contribution to axis construction. Blue vectors were the traits that contributed the least, while orange vectors were the traits that contributed the most. Culture conditions and parental strains effects were considered as covariables of traits. The size of these vectors (in black) indicates its influence on the other traits.*

Thus, subsequent analyses were conducted independently for nitrogen limited and phosphorus limited conditions. Consequently, the results of GWAS would be specific to each nutrient limitation, potentially enabling the identification of genes associated exclusively with one condition or both. Moreover, the vectors representing the principal components were examined to infer relationships between traits (Figure 1). Traits associated with lipids (TFA, SFA, MUFA, PUFA, omega3 and omega 6) were grouped and at opposite of PCA axis 2, traits associated with photosystems (Chl_a, Chl_c1, Chl_c2 and fucoxanthin) were grouped together. These two groups were the most significant contributors in the definition of the first PCA axis, highlighted two known metabolic pathways: lipid and pigment metabolisms. These correlations between these different traits were confirmed with the Spearman correlations (Sup. figure S3.7). The variations in these traits within these two groups are partly related because they belong to the same metabolic pathway (lipids or pigments) and are therefore interdependent.

In order to visualise the variations induced by the origin of the parental strains in each nutrient condition, a principal component analysis (PCA) was performed on the pigment and lipid profiles of each lineage of algae (Figure 2). The lineages were well distributed in the PCA, but they were also clustered according to their respective parental strain, thus highlighting their origin. This result was consistent with the collection algal lineages set up from parental strains, these results were not surprising and this coancestry effect was considered during the association tests.

**Figure 2.**
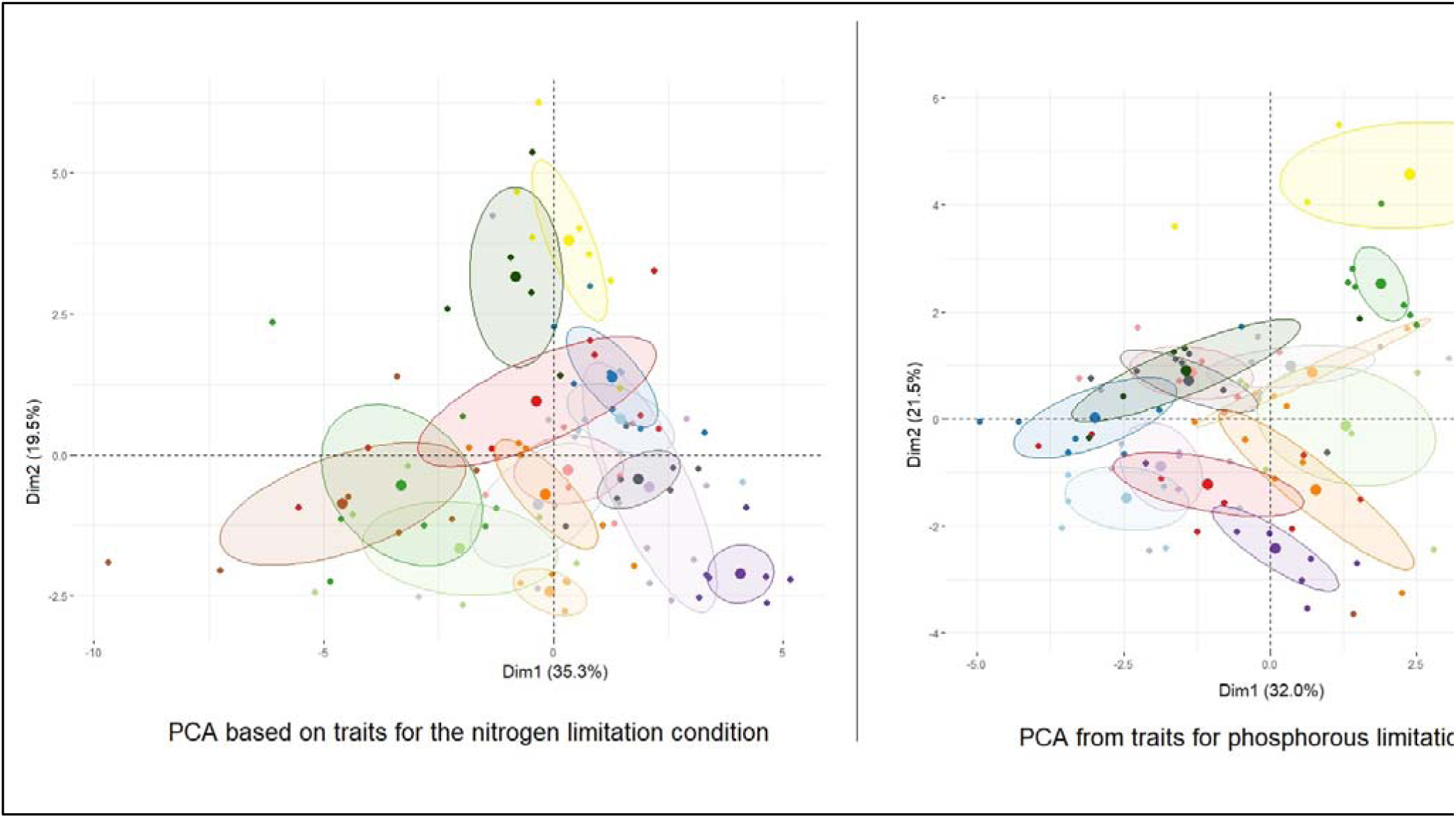
PCA based on phenotypic traits in nitrogen limitation or in phosphorous limitation. *Each dot indicates the position of a lineage in this PCA according to its phenotypic profile. Each colour represents the parental strain origin of each lineage. The ellipses represent 95% of the total variance of all lineages from a same parent strain, and similarly the large dot indicates the mean.*

### Genotypic data for GWAS

Besides phenotypic diversity, GWAS is based on genotypic diversity resulting from the plasticity of genome and the divergence of individuals (Bazakos et al., 2017). The genome of each algal lineage was thus sequenced (Sup. data S4.1). On average, the sequencing depth of the *T. lutea* genome for lineages was 132x (Sup. figure S4.2). This degree of coverage allowed the accurate identification of genetic variations between algal lineages. Then, polymorphism calling was performed between all algal lineages sequenced and 104,984 genetic polymorphisms were identified (Sup. data S4.3). These genetic polymorphisms included three types of molecular variations: 103,097 mutations as single nucleotide polymorphisms (SNPs), and short insertions or deletions (<10 bases); 804 large insertions or deletions due to transposable element (>1000 bases); and 1,083 presence or absence of genes. Compared to the previous study on parental strains (Carrier et al., 2024), the number of large variations was certainly underestimated. This was due to the use of short reads for sequencing the algal lineages. The distribution of genetic polymorphisms covered the entire genome with an average of 1.2 polymorphisms per kb. Furthermore, an average of 0.8 polymorphisms per gene was observed in their coding regions. Thus, this density of polymorphisms allowed the study of gene-phenotype associations across the entire genome.

Before GWAS analysis, it is crucial to study familial relatedness or/and population structure because they are known to increase false-positive rate (Atwell et al., 2010). Therefore, the effects of genetic ancestry and the genetic structure of the algal lineages were estimated to be taken into account in GWAS analysis (Yu et al., 2006). The genetic structure of algal lineages was inferred without prior knowledge, using likelihood-based methods (Frichot et al., 2014). The optimal ancestry coefficient was estimated at four subpopulations using the entropy measurement (Figure 3-A). A genetic similarity tree was built between algal lineages to observe their co-ancestry according to the origin of the parental strain (Figure 3-D) and the estimated coefficient of ancestry (Figure 3-C). As expected, the distribution of algal lineage in the tree was structured according to the origin of the parental strain (Figure 3-D). The subdivision into four subpopulations, defined by the coefficient of ancestry, showed very limited overlap with the parental origin of the lineages. Almost all lineages derived from four parental strains (Tiso-14, 13, 12 and 9) belonged to the same subpopulation, whereas for all other parental strains, their lineages were spread across several subpopulations. The fixation index (*F*_ST_) between these four subpopulations was weak (mean *F*_ST_ = 0.06; Sup. table S4.8.1), which suggests that the genetic structuration of this population was limited. Moreover, the fixation index measured between parental strains was also weak (*F*_ST_ = 0.04; Sup. table S4.8.2). Thus, our collection of algal lineages appeared weakly structured, with a limited parental strains effect. Interestingly, the effect of the parental strains on genetic structure was present, but weaker than expected. This suggests that evolutionary forces (such as mutation rates and drift) also favoured genetic variation among these lineages. Moreover, we performed the genetic structure analysis specifically on the chloroplast (Contig25) and mitochondrial (contig26) genomes (Figure S4.10). For the chloroplast, the results matched the whole-genome analysis, with the optimal cross entropy identified at 4 subpopulations (K=4). For the mitochondrion, the pattern was different and less clear; it was thus difficult to determine the exact optimal number of subpopulations. Regarding the number of genetic polymorphisms, their genomic distribution and the limited structuration of the population, we considered that the genotypic data were adequate to use in GWAS.

**Figure 3.**
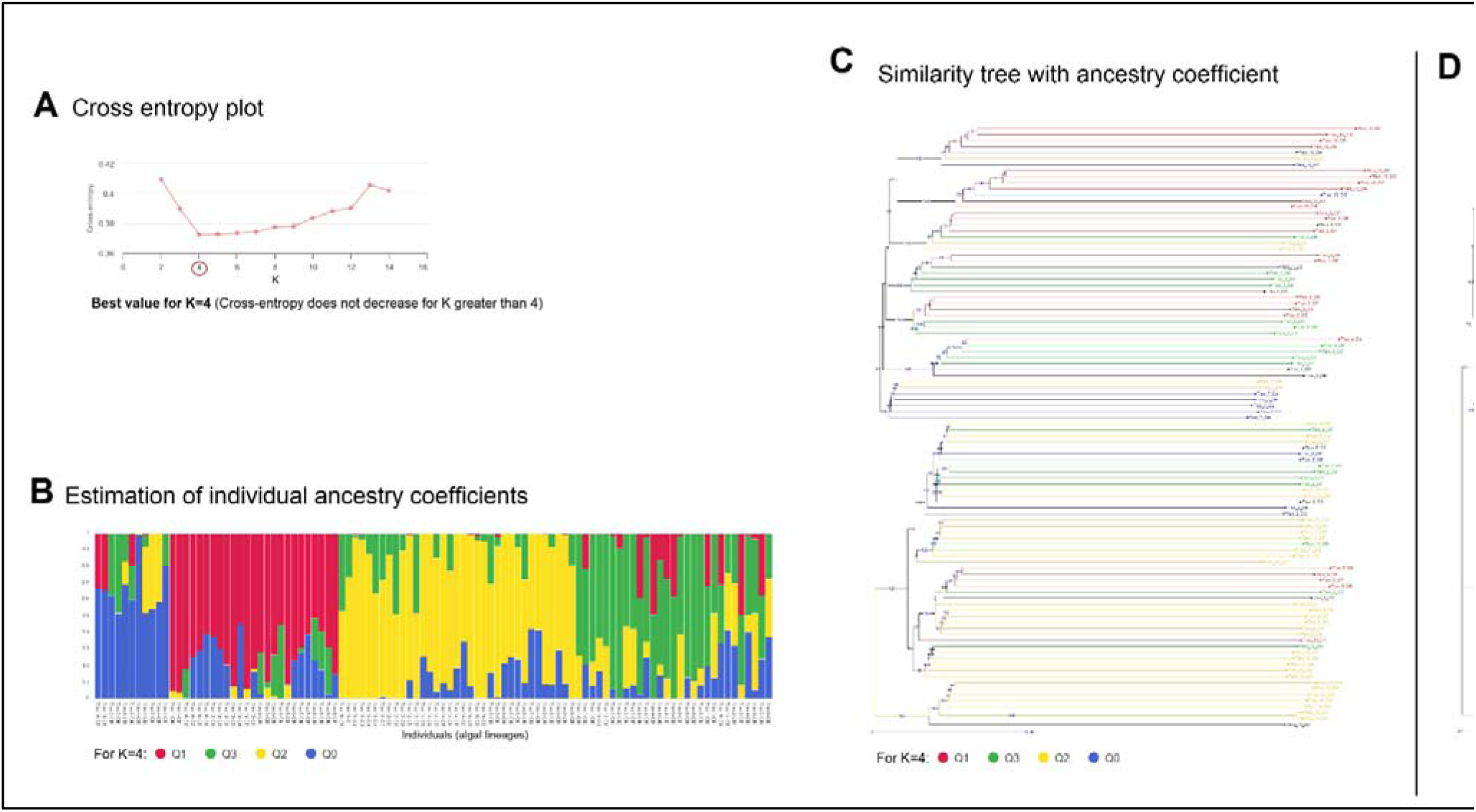
Genotype structure of different algal strains. *Figure A. Plot showing the cross-entropy score that identifies the optimal number of strain groups (K=4). Figure B. shows the genetic contribution of each of the previous four groups to each lineage. Figures C. and Figures D. show the genetic similarity tree. Figure C highlights the distribution of the four ancestral groups, and Figure D. highlights the origin of the parental strains.*

### GWAS results and characterization of associations

GWAS were conducted with the set of phenotypic and genotypic data previously described. In general, studies of quantitative traits have assumed a complex genetic architecture where traits were influenced by not just one gene, but many genes and thus several polymorphisms distributed across multiple regions (Visscher et al., 2017). Thus, an approach using multi-locus mixed models was used to consider the joint effect of several polymorphisms (Segura et al., 2012). Additionally, although the structuring of this population was limited, a kinship matrix (Sup. data S5.2) was incorporated into the association model through a polygenic term to limit false positives (Price et al., 2006; Pritchard et al., 2000). Association tests were performed using this model, and only significant associations were considered after removing those with scores that were too low (as weak contribution to phenotypic variance or failure to meet the Bonferroni threshold). Furthermore, the alignment of the observed P-values with the expected distribution across the quantile-quantile (QQ) plots (Sup. Figure S4.14) confirmed that our MLMM model effectively accounted for population stratification, thereby ensuring the statistical reliability of the detected associations.This was further confirmed by the estimation of the genomic inflation factor (λ) to evaluate the presence of false positives or residual population stratification (Yang et al., 2011). It was calculated for each trait in GWAS (Sup. Table S4.15). Across these traits and both conditions, the genomic inflation values ranged from 0.74 to 1.10, indicating an effective correction of population structure by the kinship matrix (Sup. Figure S.4.14).

Thus, this approach revealed 13 significant associations (Table 1). Among them, seven associations were linked to quantitative variation of pigments and one to non-photochemical quenching linked to global photosystem variations. The five others associations were linked to lipids variations. Each association was identified in only one culture condition (eight in nitrogen deficiency and five in phosphorus deficiency), none in both conditions. Finally, among the 13 identified associations, ten involved a nucleotide polymorphism, two an indel, and one the insertion of a transposable element. For each association, the corresponding genomic regions were determined, and initial functional hypotheses were proposed. More precisely, the score of explained phenotypic variances by the polymorphism associated, the *Fst* score, the allelic distribution, the genes around the polymorphism associated, and their putative proteins were reported (Sup. figures S5.3-S5.15). Among the 13 associations, 17 candidate genes were identified. Their proteins were analysed to propose hypotheses about their molecular function based on homologous proteins characterised in other species (Table 1). However, for six of these proteins, no homologous protein structure or protein domain was identified, meaning that these putative proteins were considered unknown. Among proteins with a putative molecular function, the link with the trait identified in GWAS appeared quite evident for some of them. For example, a polyketide synthase was identified as potentially involved in the amount of secondary pigment (C26:323312). An inorganic diphosphatase enzyme was also identified as presumed to be involved in photosynthesis metabolism (C49:2395209). Finally, a transcription factor likely affecting the amount of polyunsaturated lipids was identified (C41:1815142). For others, the link was less evident or much more indirect. Examples include a methyltransferase-like enzyme linked to chlorophyll content (C50:1371586), a hydrolase-like protein involved in diadinoxanthin content, and an aminotransferase-like enzyme affecting lipid content (C49:3135046).

**Table 1.** Significant associations identified.

| Physiological state according to nutrient deficiency | Trait associated | Position in genome | Types of genetic polymorphism | Minimum allele frequency | Association test, *P*value score | Estimation of phenotypic variance explained by a genetic polymorphism | Fst score (between parental strains) | Potentially impacted gene | Putative protein function |
| --- | --- | --- | --- | --- | --- | --- | --- | --- | --- |
| Nitrogen deficiency | Violaxanthin | Contig_31:998924 | SNP | 0.12 | 2.35E-11 | 0.55 | 0.21 | 36257 | Ubiquitin-like |
|  |  |  |  |  |  |  |  | 36259 | Unknown |
| Phosphorus deficiency | Echinenone | Contig_26:323312 | SNP and TE | 0.29 | 4.52E-09 | 0.53 | 0.26 | 37631 | Polyketide synthase |
| Phosphorus deficiency | Chlorophyle c2 | Contig_39:44927 | SNP and SSR | 0.23 | 1.03E-10 | 0.53 | 0.26 | 36259 | Elongation of very long chain fatty acids |
|  |  |  |  |  |  |  |  | 36257 | Unknown |
| Phosphorus deficiency | Non-Photochemical Quenching | Contig_49:2395209 | indel | 0.13 | 2.91E-09 | 0.46 | 0.24 | 12311 | Inorganic diphosphatase |
| Nitrogen deficiency | Chlorophyle c1 | Contig_50:1371586 | indel | 0.27 | 1.74E-09 | 0.42 | 0.44 | 15113 | Methyltransferase-like |
| Nitrogen deficiency | Echinenone | Contig_46:2354763 | SNP | 0.23 | 4.92E-11 | 0.34 | 0.25 | 21620 | Unknown |
| Nitrogen deficiency | Fucoxanthin | Contig_53:912007 | SNP | 0.32 | 3.77E-07 | 0.28 | 0.15 | 7834 | Unknown |
| Phosphorus deficiency | Monounsaturated fatty acids | Contig_39:923029 | SNP | 0.15 | 2.35E-77 | 0.2 | 0.12 | 30411 | Unknown |
|  |  |  |  |  |  |  |  | 30414 | SNF7 family protein |
| Phosphorus deficiency | Polyunsaturated fatty acids | Contig_41:1815142 | SNP | 0.28 | 6.04E-16 | 0.17 | 0.06 | 26683 | Transcription factor |
|  |  |  |  |  |  |  |  | 26887 | Transmembrane protease serine |
| Nitrogen deficiency | Diadinoxanthin | Contig_46:2397370 | SNP | 0.43 | 1.38E-07 | 0.15 | 0.19 | 21643 | hydrolase like for phospholipase |
| Nitrogen deficiency | Total fatty acids | Contig_49:3135046 | SNP | 0.46 | 3.14E-90 | 0.14 | 0.01 | 12694 | Aminotransferase-like |
| Nitrogen deficiency | Polyunsaturated fatty acids | Contig_51:3397622 | SNP | 0.21 | 1.85E-07 | 0.11 | 0.09 | 14376 | Unknown |
| Nitrogen deficiency | Polyunsaturated fatty acids | Contig_48:1297470 | SNP | 0.43 | 2.54E-24 | 0.1 | 0.03 | 16659 | Unknown |

Among all relevant genetic polymorphisms identified in this study, three of them exhibited a phenotypic variance score higher than 50%, thus attracting our particular attention (Figure 4). While such high scores highlight these loci as major-effect candidates, this proportion of explained variance may be somewhat overestimated due to residual population structure or to the low sample size (n = 100) in our dataset. Caution is therefore required when interpreting these values, and while we consider these loci to have a major impact on the phenotypic traits, the precise magnitude of their variance contribution will need to be confirmed through downstream functional experiments.

**Figure 4.**
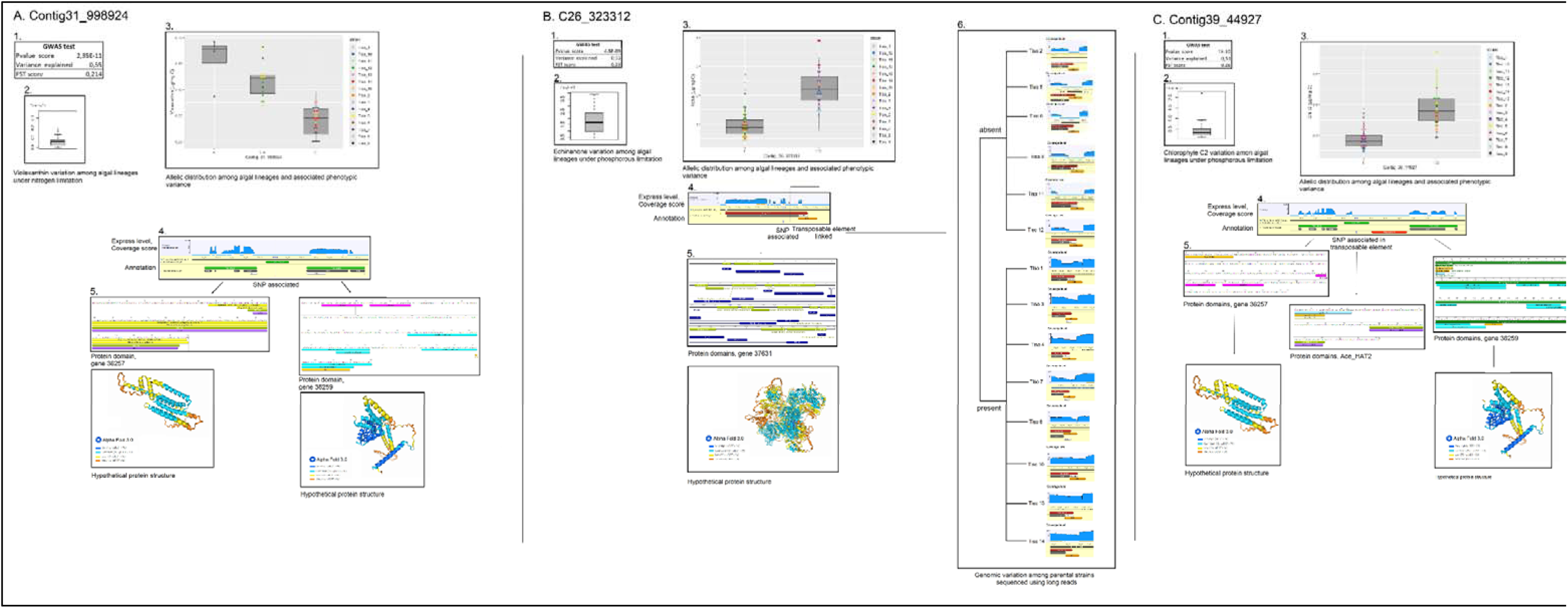
Associations with the highest explained phenotypic variance score. *For each association: 1. A table summarising the association test results, the score of phenotypic variance explained and the Fst index; 2. A box plot of phenotypic measurements; 3. Allelic distribution of loci according to phenotypic measurement; 4. Position of the loci and proximal genomic annotation; 5. Gene and putative protein which could be linked to the identified association; 6. Studies of large variations using long-read sequencing from parental strains (Carrier et al., 2024). Figure 4.A. Association between loci C31_998924 and violaxanthin content, Figure 4.B. Association between loci C26_323312 and echinenone content, and Figure 4.C. Association between loci C39_44927 and chlorophyll-c1 content.*

The C31: 998924 polymorphism obtained the highest explained variance score in relation to violaxanthin pigment content (Figure 4-A). Homozygous algal lineages (A) exhibited high violaxanthin content under nitrogen-depleted conditions. In contrast, homozygous lineages (C) displayed significantly reduced violaxanthin content, and lineages with both alleles (A:C) showed an intermediate level (Figure 4-A.2). Therefore, selection of homozygous lineages (A) should yield lineages with higher violaxanthin under nitrogen-depleted conditions. Moreover, the mean *Fst* index between parental strains was high (0.21) whereas for almost the entire genome it was only 0.04. The parental strains Tiso-4, 5 and 12 carried this allele (A). Consequently, this genetic polymorphism could have been selected because it provided an adaptive advantage to several parental strains in their environment over others. Unfortunately, the information about their geographical origins was too ambiguous to be interpreted. This genetic polymorphism was located in an intergenic area between two genes (36257 and 36259), but its direct impact on these genes was not evident (Figure 4-A.3). The gene 36257 contained protein domains associated with a ubiquitin-like protein. This protein is mainly found in haptophytes (Sup. figures S5.15). Furthermore, analyses of the putative proteins did not show a clear connection with secondary metabolism and requires further studies to formulate hypotheses about their own biological function.

The second polymorphism, C26: 323312, was associated with echinenone pigment content (Figure 4-B). Lineages that possessed both alleles (T/C) exhibited a higher echinenone content than lineages that possessed only the (T) allele. However, none of the studied lineages possessed only the (C) allele. It was likely that, in its homozygous form, this allele could have been deleterious to the alga. The associated polymorphism was located in the coding region of the gene, but appeared to have limited impact on the protein because it did not change the amino acid sequence. However, manual analysis revealed the presence of another genetic polymorphism due to a large insertion/deletion near the associated SNP. A sequence similarity search indicates that this genetic polymorphism may be the result of an LTR-type transposable element from the LAR family being inserted (Berthelier et al., 2018). Indeed, short-read sequencing inherently limits the accurate detection and characterization of such structural variants, creating a degree of uncertainty when interpreting complex genomic loci. Unfortunately, this genetic polymorphism was not included in the initial analysis because its size (around 1,000 pb) was insufficient to be considered as a potentially large variation. Yet, this large insertion/deletion was in complete linkage disequilibrium (D’ = 1) with the previous significantly associated polymorphism (C26: 323312) and therefore it can be considered as similarly associated. Moreover, the *Fst* index between the parental strains was high (0.22), suggesting positive selection between two groups of parental strains ((Tiso-2, 5, 6, 9, 11, 12) and (Tiso-1, 3, 4, 7, 8, 10, 13, 14)). To overcome the limitations of short-read data, long read sequences analysis of the parental strains (Carrier et al., 2024) was used and confirmed the observed polymorphisms and the structuring into two groups of strains (Figure 4-B.6). Parental strains lacking the large insertion/deletion (Tiso-2, 5, 6, 9, 11, 12) exhibited the highest echinenone content. In contrast, parental strains harboring an insertion showed the lowest echinenone levels under phosphorus-depleted conditions (Tiso-1, 3, 4, 7, 8, 10, 13, 14). These results strongly suggest that this insertion impacts the neighboring gene 37631. Ideally, to fully capture the impact of structural variations, future GWAS should employ long-read genotyping across all studied lineages. The putative protein associated with this gene was identified as a polyketide synthase-type (PKS) protein. Members of this protein family are known to catalyse the biosynthesis of secondary metabolites structurally related to the echinenone pigment. Proteins homologous to PKS were identified in a wide range of marine taxa including Haptophyta, Chromista, Chlorophyta and Opisthokonta (Sup. figure S5.15). However, PKS are functionally diverse, which limits the possibility of specifically inferring their biological roles solely from sequence similarity. Consequently, further investigation of the molecular function of this enzyme is warranted and is particularly relevant for understanding the biosynthesis of this secondary pigment.

The genetic polymorphism, C39_44927, appears to be associated with a content of chlorophyll *c2* (Figure 4-C). This polymorphism is located in a transposable element annotated (Ace_Hat2), and near to a simple sequence repeat (SSR). The Ace_Hat2 element was previously identified as potentially active (Berthelier et al., 2023). In a first analysis using short-reads sequencing, this insertion appeared to be present in all lineages. In parental strains sequenced using long-reads, the genotyping of this insertion remained ambiguous. Indeed, in the parental strains Tiso-2, 5, 6, 7, 9, 11, and 12, a coverage bias, corresponding to an accumulation of reads at TE insertions was observed. This biased coverage could have been interpreted as a mapping artefact due to identical reads generated by a high activity of the Ace_Hat2 element in these strains. Alternatively, it could have been interpreted as true allelic polymorphism, meaning that these strains would have several alleles (with or without insertions) compared to other parental strains. Variations between strains was true, but it was hard to draw more conclusions based on this data set. Analysis of the SSR was not easier, especially when they are large, as it is the case here (larger than 1,000 bases). Short read sequencing was therefore not sufficiently informative for genotyping algal lineages. Long reads sequencing of the previously sequenced parental strains showed strong variation of this SSR (Figure 4-C.6). Parental strains Tiso-1, 13 and 14 exhibited the presence of the large motif of this SSR, while in other parental strains, this motif was considerably reduced. The polymorphism of this SSR was established in parental strains, but unfortunately, based solely on these data, it was difficult to characterise this SSR in relation to the associated loci. Globally, it was not easy to describe this hotspot of genetic variation, but polymorphism between strains was undeniably present. In the future, a depth-sequencing approach involving long-read sequencing in each lineage, should be employed to decipher genetic polymorphisms and link them to their respective phenotypes. Two genes (36257 and 36259) were present around this association revealed with GWAS. Variation in the regulation of these genes could explain the observed phenotypic differences. The protein putatively encoded by gene 36257 had neither recognizable domain or homologous structure, making any further interpretation difficult. The second gene 36259, could encode an elongase enzyme involved in the synthesis of very long-chain fatty acids. This elongase could play a role in the synthesis of phytol chains in chlorophyll *c2* or MGDG (monogalactosyldiacylglycerol) fatty acid chains in thylakoid membranes which were known to interact in this species (Gonçalves de Oliveira-Júnior et al., 2020). Homologs of this protein were identified only in haptophyte organisms, suggesting a degree of specificity for this taxon (Sup. Figure C-S5.15). Additional studies are needed in the future to validate these hypotheses. Of the three genetic polymorphisms highlighted, this locus will require the most in-depth investigation to elucidate its molecular implication.

## General discussion

The number of genes inventoried for all oceanic organisms was estimated around 1.5 million (Carradec et al., 2018). However, only about a quarter of them have enough similarities to genes or proteins identified in other organisms to be assigned a phenotypic function. Furthermore, the genes that were most easily assigned to characterised homologues were those that are best conserved in living organisms and therefore belonging mainly to central metabolism (Koonin, 2005). Regarding other phenotypic traits, few genes were described, particularly in marine organisms. Using quantitative genetics to propose functional hypotheses for genes without a priori has already proven effective in many model organisms, especially humans. Indeed, the NHGRI-EBI GWAS Catalog currently contains over 5,000 GWAS studies on humans. (Sollis et al., 2023). Numerous GWAS studies have also been conducted in the agronomic field to search for genetic markers in the context of breeding programmes (Alseekh et al., 2021). However, in the context of marine organisms, the number of studies is extremely limited. Only a few studies have been conducted in oysters (Liu et al., 2022) and a few farmed fish such as sea bass (Oikonomou et al., 2022). The objective of this study was to perform GWAS in phytoplankton to better understand the role of genes by identification of genetic loci linked to phenotypic traits. In light of the results obtained in this study, the limitations and issues for improvement will be discussed to determine whether this approach could be effective in a marine context.

### Biological resources, the major challenge

GWAS requires a collection of several dozen (or even hundreds) of individuals (Ott et al., 2011). Exploiting only 100 individuals is a relatively modest sample size for GWAS today given the high marker density, but working with microalgal collections presents logistical constraints. Unfortunately, the harvesting, isolation, and preservation of living phytoplankton were not yet performed on a large scale. Collecting one hundred life samples through marine campaigns is very time-consuming and complex. Currently, most marine sampling campaigns for biodiversity inventories are unable to conserve each specimen in a living collection. The preservation of marine organisms is far more complex and costly than that of terrestrial species due to the challenges posed by the marine environment. This is currently the main obstacle to implementing quantitative approaches. Concerning the microalga *T. lutea*, only 15 parental strains from different oceans were available in collections, which was insufficient for implementing GWAS approaches. To partially address this problem, we exploited the individual diversity present within each of the strains (Carrier et al., 2024). Microalgal strains were not composed of pure lineages but encompassed a broad range of genotypes, with each strain representing a subpopulation of the species (Lakeman et al., 2009; Sjöqvist, 2022). The existence of individual diversity within strains of this species has previously been acknowledged and has subsequently been employed in breeding programmes (Gachelin et al., 2021). Nevertheless, the biological processes driving this intra-diversity remain unclear. The mutation rate and the effect of genetic drift was not yet quantified in this species. In few cases, the mutation rate may contribute to this intra-diversity (Krasovec et al., 2018, 2017; Ness et al., 2012). The exploitation of intra-diversity from the 15 parental strains of *T. lutea* enabled the generation of 100 distinct algal lineages. The biological dataset utilized in this study was not optimal, as it does not represent the full biological diversity of the species. Obtaining 100 lineages from only 15 parental strains intrinsically generates a hierarchical population structure. As expected, phenotypic analyses revealed significant structuring among lineages in function on strain origin, as evidenced by high Q_ST_ values (0.5) and PCA clustering. However, genetic metrics, such as F_ST_ scores (0.04) indicate a relatively weak genetic structuring within this population. This stark contrast in population structure between the phenotypic and genetic levels is surprising given the hierarchical design of our population. Although the presence of high intra-strain genetic diversity has been previously documented (Carrier et al., 2024), we anticipated a higher genetic stratification. Q_ST_ values higher than F_ST_ scores could suggest the presence of mechanisms promoting divergence among individuals within strains. Nevertheless, because the life cycle of this species remains largely uncharacterized, formulating robust evolutionary hypotheses is currently challenging. Elucidating the life cycle of this microalga is therefore a crucial prerequisite for future studies aiming to fully understand these population dynamics.

### Phenotyping under varied environmental conditions

In this study, cultures were grown for each lineage in duplicate and under two nutritive conditions (nitrogen or phosphorus starvation). The analysis showed that the environmental effect strongly influenced phenotypic variation. Therefore, each culture condition was analysed independently. Consequently, we were not able to use methods such as BLUPs (Speed and Balding, 2014) to subtract the portion of variance not attributable to the environment. This implies that the genetic associations identified in this study are dependent on specific environmental conditions. Because these associations are strictly environment-specific, the generality of our findings is inherently limited; the identified candidate genes may only drive phenotypic variation under these specific nutrient starvation conditions rather than under standard or alternative environments. Furthermore, the current experimental design with only two conditions did not allow for the integration of Genotype-by-Environment (GxE) interactions or the estimation of environment-independent genetic effects, which restricts our ability to disentangle constitutive genetic drivers from purely stress-responsive alleles. Moreover, in marine ecosystems, nutrient limitation is the prevailing physiological state, whereas nutrient-replete conditions are highly transient exceptions (Moore et al., 2013). Consequently, rather than employing an artificial, nutrient-saturated control, our experimental design deliberately focused on nitrogen and phosphorus limitations. These conditions represent the most ecologically relevant constraints for marine microalgae, which have evolved to function primarily in chronically depleted environments. While nitrogen and phosphorus availability dictate the primary metabolic strategies of these organisms in the ocean, expanding this framework to include a nutrient-replete baseline and other environmental drivers like light and temperature remains an important perspective. In future GWAS approaches, it will be necessary to significantly increase the number of environmental conditions studied and formally integrate GxE models, as is now more frequently the case for other species such as Arabidopsis and Drosophila (Burke et al., 2014; Morgante et al., 2015).

### GWAS results

GWAS was performed using a Multi-Locus Mixed Model (MLMM) pipeline combined with a kinship matrix to maximize statistical power and limit false negatives while controlling for population structure. QQ-plots and genomic inflation factors confirmed the robustness of the MLMM utilizing the kinship matrix, showing adequate control of population stratification. Nonetheless, caution is warranted with a high phenotypic structuring. There remains a residual risk that some of the detected associations might reflect shared ancestry or cryptic population stratification rather than true causal genotype–phenotype relationships. Consequently, while our strategy successfully circumvents the biological bottleneck of marine strain availability, the identified loci must be interpreted cautiously as primary candidate regions that will ultimately require downstream functional validation. In total, the GWAS approach identified 13 associations between genetic polymorphisms and phenotypic traits. In other words, the identified allelic polymorphisms contributed, at least in part, to the variability of the phenotypic trait. To understand the genetic basis of these associations, the hypothesis was formulated that the proximal genes to an associated genetic polymorphism can be considered involved in the variation of the phenotypic trait. This gene could affect the phenotypic trait directly or indirectly, and positively or negatively. Yet, GWAS results must be interpreted with caution, as the relationship between a genomic position, the proximal gene and the trait can be very complex. The associated genetic polymorphism may influence the expression of a gene or other component of the genome located far away, (Maurano et al., 2012; Visscher et al., 2017). Among the 16 identified associations in this study, three primary candidate loci emerged as top priorities, exhibiting a high proportion of explained phenotypic variance and high statistical association. Furthermore, the homologous functions of their proximal genes are consistent with the associated phenotypes. Our hypotheses include: an intergenic region linked to a ubiquitin-like domain protein (gene 36257) associated with violaxanthin levels, a transposable element insertion impacting a polyketide synthase (PKS, gene 37631) potentially involved in echinenone biosynthesis, and a complex locus harboring a fatty acid elongase (gene 36259) putatively linked to chlorophyll *c2* and thylakoid lipid regulation. While these candidate genes offer highly compelling mechanistic leads, these functional roles remain very speculative at this stage. Consequently, targeted functional characterization will be clearly necessary to understand and confirm the molecular mechanisms and their implications in bioprocesses.

## Conclusion

This GWAS approach allowed us to identify candidate loci associated with the traits of interest in *Tisochrysis lutea*, an outcome that would have been highly challenging to achieve using alternative methodologies. Currently, there is no straightforward method to fully characterize these genomes. In addition, applying functional approaches, such as CRISPR, to marine species continues to present a considerable challenge. Indeed, despite ongoing efforts in our laboratory to establish a reliable genetic transformation protocol for this alga, successful implementation remains a significant hurdle. A major advantage of GWAS is its ability to identify candidate genes without *a priori* knowledge. In our study, it would have been impossible to deduce the involvement of these loci in the associated traits based only on genome annotation or other available data. Nevertheless, implementing this approach required significant investment to identify a dozen associations, and the greatest challenge in the marine field remains the establishment of large-scale strain collections. In this specific microalgal context, intra-strain diversity allowed us to overcome the difficulty of accessing a wide range of strains. However, in the future, it will be necessary to obtain new strains from diverse environments and increase phenotyping across various environmental conditions to optimize the efficiency and statistical power of the GWAS approach. Regardless of the methodology employed, a substantial effort from the scientific community is required to better understand marine genomes. In this context, quantitative genetics remains a highly valuable strategy for advancing our understanding of the genetics of marine organisms.

## Materials and methods

### Collection of algal lineages

In this study, we established a collection of 15 T. lutea algal lineages from a worldwide collection (Sup. Table S1.16). These 15 strains were isolated and considered as parental strains. They were not pure lineages, but rather populations exhibiting inter-individual diversity (Carrier et al., 2024). The 15 T. lutea strains studied were: CCAP927-14, CCMP463, RCC179, RCC1344, RCC3691, RCC3692, RCC3693, RCC3699, NIVA 4-91, IMFG, Argenton, S2M2, MCCV95, MCC97, and S5. Parental strains were maintained in 200 mL flasks with natural seawater enriched with sterile Conway-Walne medium (Walne, 1966), and transferred to fresh medium every three weeks. Cultures were maintained at a constant temperature of 21°C, under a constant light irradiance of 50 μmol m−2.s −1 and bubbled with 0.22 µm filtered-air.

To obtain algal lineages, individual cells were isolated using a flow cytometer based on their lipid and pigment content. Prior to the sorting process, each parental strain was cultivated under identical conditions for six days under nitrogen deficiency. Sorting was performed using a FACS Aria II Cytometer at the Cytocell platform (Nantes, France). Chlorophyll fluorescence was directly measured using a PerCP-Cy5.5 filter, and neutral lipid content was indirectly measured using a Nile red fluorochrome (Cas. No: 7385-67-3)) (Rumin et al., 2015). The excitation wavelength used for Nile red was λ=525 nm and the emission wavelength was λ=580 nm. Each new algal lineage was isolated within a specific gating window based on Nile red and chlorophyll fluorescence measurements (Sup. figures S1.1 to S1.15). The objective was to obtain algal lineages with diverse lipid and pigment profiles. For each sorted parental strain, 96 lineages were generated by isolating single cells into 200 μL of culture medium in sterile 96-well plates. After approximately 45 days of culture, involving a gradual step-up in culture volume, a sufficient biomass was reached (10 million of cells. mL-1 in 100mL). Between 0 and 73 lineages survived for each of the selected sorting gates (Sup. table S1.17). Finally, one algal lineage was randomly selected per gate to represent each parental strain, resulting in a total collection of 100 algal lineages (Sup. table S1.17). Algal cells were re-suspended for conservation every three weeks.

### Culture apparatus and experiment conditions

To facilitate the gene-phenotype association approach, cultures were grown in a custom “phenotyping bench” device (Berard et al., 2021) specifically designed to minimize environmental variability within and between experimental runs (Sup. data S2.1.1). The apparatus consisted of 24 photobioreactors with precise regulation of temperature, pH, airflow, and irradiance. By systematically standardizing these parameters, all experiments were performed under highly homogeneous and reproducible conditions. Based on this approach, each algal lineage was grown in biological duplicates in batch mode under two nutrient-limited conditions: nitrogen-limited (Nlim) and phosphorus-limited (Plim) (Sup. data S2.1.1). To ensure equitable growth initiation across all algal lineages, the pre-acclimation of the inoculum and the initial cell concentration were standardized to 300 000 cells.mL^-1^ for each reactor. For each algal lineage, growth conditions were strictly controlled (constant irradiance of 350 μmol m^−2^·s^−1^, temperature 23°C, pH 7.5, airflow 4 L.h^-1^) using Conway nutrient medium (Walne, 1966) modified with 150 µM NO_3_ for nitrogen limitation and 5 µM of PO_4_ for phosphorus limitation. Biomass growth was automatically monitored via optical density measurements (Sup. data S2.3.0 to S2.3.21). Manual optical density measurements were performed daily using spectroscopy at 680 nm. The depletion of the limiting nutrient (nitrogen or phosphorus) was also monitored spectroscopically. Microalgal cells were removed by filtration (0.2 µm), and the culture medium was analyzed as described by Collos et al., (1999) for nitrogen and (Van Veldhoven and Mannaerts, 1987) for phosphorus.

### Phenotyping algal lineages

Each algal lineage was cultivated under identical conditions in biological duplicates (Sup. figures S2.1.2 and S2.1.3). The cultivation process began with a growth phase, during which the nutrients in the medium were consumed (approximately two days). Upon depletion of the limiting nutrient, microalgal growth continued by utilizing intracellular reserves (for approximately two additional days). Once these reserves were depleted, the culture entered a stationary phase. Harvesting for phenotyping was carried out two days into the stationary phase, corresponding to day six or seven of the culture. During the harvest of each algal lineage, multiple samples were collected for phenotyping measurements. Approximately 8 mL of culture (about 25 million algal cells) was filtered through pre-combusted GF/C filters (Whatman, 25 mm diameter) to separate the microalgae from the medium. Filters were processed according to the specific downstream analysis: i) flash-frozen in liquid nitrogen and stored at -80°C for pigment quantification; ii) placed in 6 mL of Folch’s solution (chloroform-methanol, 2:1, v/v) and stored at -80°C for lipid profiling; iii) dried in an oven for cellular organic carbon measurement. Simultaneously, 2 mL aliquots of the algal culture were immediately sampled to determine exact cell concentration and size, and an additional 2 mL was collected for active fluorescence measurements.

#### Cells number and size

Prior to measurement, algal culture samples were diluted 1:50 with sterile seawater. Cell size and number were assessed using a Coulter Counter Multisizer 3 (Beckman Coulter, High Wycombe, UK). Cell size, given as equivalent spherical diameter, was calculated using the MS-Multisizer 3 software.

#### Measures of organic carbon in algal cells

Filters containing the harvested algal biomass were placed in glass vials and dried in an oven at 75°C for 24 h. Carbon content was measured using a carbon elemental analyzer (Thermo Electron), calibrated with methionine, aspartic acid, and nicotinamide standards.

#### Measures of lipid profile

The extracts containing fatty acids were evaporated and converted into fatty acid methyl esters (FAMEs) by acid-catalyzed transesterification with methanol, according to the “FAME Preparation” procedure described in Guihéneuf et al., (2015). The upper organic phases containing FAMEs were collected and assayed by separation on a ZEBRON™ ZB-FAME column (30 m × 0.25 mm × 0.20 μm, Phenomenex™) using gas chromatography (TRACE™ 1310, Thermo Scientific™) coupled with a flame-ionization detector (GC-FID). FAME quantification was performed using methyl heptadecanoate (C17:0) as an internal standard (Sigma-Aldrich, St. Louis, MO), while the qualitative standard for FAMEs was the Supelco™ 37-Component FAME Mix (Sigma-Aldrich). Total fatty acids (TFAs) were calculated as the sum of saturated fatty acids (SFAs), polyunsaturated fatty acids (PUFAs), including docosahexaenoic acid (DHA) and monounsaturated fatty acids (MUFAs).

#### Measures of pigment profile

Pigments were extracted using 2 mL of cold aqueous acetone (95%) containing 2.5 ng.L^-1^ of vitamin E as an internal standard. The mixture was vortexed, sonicated for 10 min, and incubated for 24 h at -20°C in the dark, followed by filtration through a 0.2 µm PTFE filter (Phenomenex, France). The filtered extract was analyzed by HPLC (Agilent LC 1200) with a DAD detector at 436 and 450 nm. Chromatographic conditions were as described in Pajot et al. (2023). Quantification was performed using external calibration against pigment standards (DHI Lab products, Denmark): Chlorophyll *c2* (Chl *c2*); Fucoxanthin (Fx); Chlorophyll *a* (Chl *a*); Diadinoxanthin (Dd); Diatoxanthin (Dt); Zeaxanthin (Zx); Echinenone (Echin); Violaxanthin (Vx); Pheophytin *a* (Pheo *a*); and β-carotene (β-Car). Chlorophyll *c1* (Chl *c1*), 3-hydroxyechinenone (HEchin), and cis-fucoxanthin (Cis-Fuco) were quantified using Chl *c2*, Echin, and Fx standards, respectively.

#### Photosynthetic performances: Active Chl a fluorescence measurement

Active chlorophyll *a* fluorescence was measured using a Phyto-PAM (Pulse Amplitude Modulated) Fluorometer (Walz GmbH, Effeltrich, Germany). The fluorometer applied a saturating pulse to the incubated sample and measured a fluorescence induction curve (detected at 680 nm), which was used to estimate the minimum fluorescence (F0 if dark-acclimated), the steady-state fluorescence in light FS), and the maximum fluorescence Fm if dark-acclimated and Fm’ if light-acclimated). We estimated the apparent maximum quantum yield of PSII after 2 min (Fv/Fm2min) and after 15 min (Fv/Fm15min) of dark acclimation as described by Rosenqvist and van Kooten, (2003).

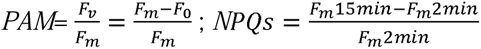

### Statistical and analyses of phenotypic data

All data analyses were performed using R (version 4.0.4). Variations in phenotypic traits were evaluated using descriptive statistics (mean, median, variance, standard deviation), and particularly boxplots and the interquartile range (IQR) (Sup. figure S3.2). The distribution of each phenotypic trait was assessed by plotting histograms and performing Shapiro-Wilk and Kolmogorov-Smirnov tests (Sup. figures S3.3 and S3.4). Traits were considered sufficiently variable if their IQR was above 0.40, and were assessed for normality based on test p-values. Phenotypic variance components were evaluated by constructing a mixed-effects model that included both fixed environmental effects and the hierarchical structure of the genetic lineages. Algal lineages were considered as nested random effects within parental strains, and parental strains as random effects within culture conditions

The model was structured as follows:

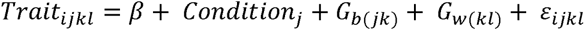

- β represents the global intercept;
- Condition (Nlim or Plim) was treated as a fixed effect to evaluate the average response across the experimental treatment;
- G_b(jk)_ represents the random effect of the interaction between condition and lineage (G x E). It accounts for the deviation of the *k*-th lineage’s response from the mean under the *j*-th condition;
- G_w(kl)_ represents the hierarchical random effect of the *k*-th lineage nested within the *l*-th parental strain. G_w(kl)_ partitions the variance attributable to the ancestral background and the specific diversification of lineages within those strains;
- □_ijkl_ is the residual error, assumed to be normally distributed From this model, we estimated the genetic component for for phenotypic traits, the heritability (*H²*) as defined by Nyquist & Baker (1991):

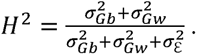

To estimate parental strains effect, the Q*_ST_* indices, an equivalent to the F*_ST_*differentiation index for quantitative traits (1993) was used: 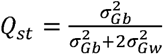.

To estimate differences between culture conditions and parental strain an analysis of variance (ANOVA) with an alpha threshold of 0.05 was performed using a general linear model (GLM). Principal component analyses (PCA) were performed on pigment and lipid contents to explore variations among algal lineages. First, PCAs were performed on pigment and lipid content in algal lineages with a nutrient limitation and parental strains considered as supplementary variables to not influence axis construction (Figure 1). Then, PCAs were performed on algal lineages separately for pigment and lipid content (Figure 2).

### Genotyping algal lineages

#### Genome sequencing

The whole genome of each algal lineage was sequenced from cultures grown under the previously described standard conditions for six days, just before their phenotyping (Conway-Walne medium, 21 °C, constant light irradiance of 50 μmol m− 2.s − 1 and bubbled with 0.22 mm filtered-air). Approximately 250 million cells (50 mL) were centrifuged (1000g), and the algal pellet was stored at -80°C. DNA extractions were conducted using the phenol-chloroform method described by Hu et al., (2004). Sequencing was performed in paired-end mode (2x150 bp) at the GenoToul platform (Toulouse, France) using Illumina NovaSeq technology. Raw data were filtered with TrimGalore (https://github.com/FelixKrueger/TrimGalore) to remove Illumina residual adapters and artifact sequences, retaining only paired reads with a quality score higher than Q30 over 150 bases (Sup. table S4.1).

#### Polymorphism identification

SNPs and short indels were identified among all algal lineages. Reads were mapped to the *T. lutea* reference genome (Carrier et al., 2024) using BWA and Mosaik software independently (Lee et al., 2014; Li and Durbin, 2009). Detection of SNPs and short indels was conducted with Freebayes and GATK independently (Garrison and Marth, 2012; Poplin et al., 2018) (Sup. data S4.4). Large structural variations, including the presence or absence of transposable elements and genes among algal lineages, were also identified. Reads of each algal lineage were mapped on the anchor genome built previously (Carrier et al., 2024). Reads were counted for each algal lineage and a presence of TE was considered depending on the number of mapped reads (minimum coverage of 50 % with 20 minimal reads depth). In the same way, genes presence/absence among algal lineages were identified (Sup. data S4.4). The polymorphism dataset was filtered using R (version 4.0.4) to remove polymorphisms with missing data higher than 5% or a minor allele frequency (MAF) lower than 10%.

### Statistical analyses of genotype data

A genetic similarity tree of the algal lineages was constructed to assess kinship using FastME 2.0 with the BIONeighborJoining distance algorithm and 1000 permutations (Lefort et al., 2015). The origin of algal lineage in function of parental strains was plotted on the tree (Sup. data S4.6).

Individual ancestry coefficients were estimated based on sNMF algorithms (Sup. data S4.7) (Frichot et al., 2014). The fixation index (F_ST_) (Weir and Cockerham, 1984) was measured for each genetic polymorphism using VCFtools (Danecek et al., 2011) between pairwise for each of parental strains and for each ancestry coefficients (Sup. data S4.8). The kinship matrix was computed from genetic polymorphisms among algal lineages as described by VanRaden, (2008). To evaluate the presence of false positives or population stratification, the genomic inflation factor λwas calculated for each trait in GWAS under nitrogen (N) and phosphorus (P) limitations based on the median of the resulting χ2 statistics (see Supplementary in Script GWAS_with GenomicInflation.R).

### Genome-wide association tests

Finally, genetic association analysis was conducted using a Multi-Locus Mixed-Model (MLMM) approach (Segura et al., 2012). Prior to the analysis, genetic variants were filtered for a minor allele frequency (MAF > 5%) and all phenotypic values were standardized using a Z-score transformation. The mixed model incorporated a random polygenic effect, utilizing a kinship matrix as the covariance structure to control for population stratification. The MLMM algorithm was parameterized with a maximum of 100 forward-inclusion steps. To minimize false positive associations, the optimal model was selected based on the multiple-Bonferroni criterion (Segura et al., 2012), applying a strict significance threshold (P < 4.76E10-7). In addition, only significant associations explaining at least 10% of the phenotypic variation were retained (Zdobnov and Apweiler, 2001). Finally, the robustness of the model for each trait was evaluated using quantile-quantile (QQ) plots (Figure S4.14) and genomic inflation factors (see Supplementary in Script GWAS_with GenomicInflation.R).

### Characterization of genes subjacent to GWAS-associations

For each identified association, the locus was characterized using available data (Sup. data S5.3 to S5.14). The F_ST_ index, the proportion of explained variance, and the allelic distribution were plotted. The genomic regions surrounding the associated polymorphisms (within approximately 1000 bases) were analyzed to identify potentially impacted genes. For each candidate gene, expression levels were plotted from available transcriptomic data (Pajot et al., 2023). Putative proteins were analyzed for functional domains using InterProScan (across all databases) (Zdobnov and Apweiler, 2001), and hypothetical structures were defined using AlphaFold (Jumper et al., 2021), followed by homologous function searches with Foldseek (van Kempen et al., 2024). Predicted effects of mutations in coding regions were estimated with SnpEff (Cingolani et al., 2012). For variations induced by a mobile element (Figure 4), confirmation was performed based on long read sequencing available for parental strains (Carrier et al., 2024). For the most promising proteins, a search for homologues in marine organisms was carried out using the Blast program (Pvalue < E-10) on Tara Ocean data (SMAGs). A krona figure (Sup. data S15.5) representing taxonomic distribution of homologs was performed by Ocean Gene Atlas web tools.

## Acknowledgements

We would like to thank SEBIMER platform for its assistance in data management and use of the Datarmor supercomputer; The Cytocell platform for sorter cells and GenoToul platform for sequencing algal lineages.

## Author contribution

Grégory Carrier supervised the research project. Thomas Lacour, Jean-Baptiste Berard, Aurelie Charrier, Elise Robert, Nathalie Schreiber, Bruno Saint-Jean, and Grégory Carrier performed the cultures and phenotyping of the different algal lineages. Vincent Segura, Agathe Maupetit, and Grégory Carrier conducted the GWAS statistical analyses. Elodie Nicolau, Anne Pajot, Bruno Saint-Jean, and Gael Bougaran provided their expertise on microalgal physiology, as well as lipid and pigment metabolism. All authors reviewed and approved the final manuscript.

## Funding for this study

This study is funded by the French National Research Agency, through the DynAlgue (JCJC) 2017-2022 project, and the French Research Institute for Exploitation of the Sea (Ifremer).

## Data availability

Method and supplementary data were available here: https://gitlab.ifremer.fr/gc0ca62/dynalgue-project.

The raw sequencing data are available in the European Nucleotide Archive (ENA) under BioProject accession number PRJEB108760 (https://www.ebi.ac.uk/ena/browser/view/PRJEB108760).

